# Directed biomechanical compressive forces enhance fusion efficiency in model placental trophoblast cultures

**DOI:** 10.1101/2024.02.22.581638

**Authors:** Prabu Karthick Parameshwar, Chen Li, Kaline Arnauts, Junqing Jiang, Sabra Rostami, Benjamin E. Campbell, Hongyan Lu, Derek Hadar Rosenzweig, Cathy Vaillancourt, Christopher Moraes

**Affiliations:** Department of Biological and Biomedical Engineering, McGill University, Montréal, Québec, Canada; Department of Chemical Engineering, McGill University, Montréal, Québec, Canada; Department of Surgery, McGill University, Montréal, Québec, Canada; Injury, Repair and Recovery Program, Research Institute of the McGill University Health Centre, Montréal, Québec, Canada; Institut National de la Recherche Scientifique (INRS)-Centre Armand-Frappier Santé Biotechnologie, Laval, Québec, Canada; Department of Obstetrics and Gynecology, Université de Montréal, and Research Center Centre Intégré Universitaire de Santé et de Services Sociaux (CIUSSS) du Nord-de-l’Île-de-Montréal, Montréal, Québec, Canada; Goodman Cancer Research Centre, McGill University, Montréal, Québec, Canada; Division of Experimental Medicine, McGill University, Montréal, Québec, Canada

**Keywords:** Placenta, choriocarcinoma, fusion, mechanics, compression

## Abstract

The syncytiotrophoblast is a multinucleated structure that arises from fusion of mononucleated cytotrophoblasts, to sheath the placental villi and regulate transport across the maternal-fetal interface. Here, we ask whether the dynamic mechanical forces that must arise during villous development might influence fusion, and explore this question using in vitro choriocarcinoma trophoblast models. We demonstrate that mechanical stress patterns arise around sites of localized fusion in cell monolayers, in patterns that match computational predictions of villous morphogenesis. We then externally apply these mechanical stress patterns to cell monolayers and demonstrate that equibiaxial compressive stresses (but not uniaxial or equibiaxial tensile stresses) enhance expression of the syndecan-1 marker of fusion. These findings suggest that the mechanical stresses that contribute towards sculpting the placental villi may also impact fusion in the developing tissue. We then extend this concept towards 3D cultures and demonstrate that fusion can be enhanced by applying low isometric compressive stresses to spheroid models, even in the absence of an inducing agent. These results indicate that mechanical stimulation is a potent activator of cellular fusion, suggesting novel avenues to improve experimental reproductive modelling, placental tissue engineering, and understanding disorders of pregnancy development.

## Introduction

The placenta is a dynamic organ that serves as the main interface between the mother and the developing fetus^1,2^. The chorionic placental villus is the fundamental unit of this interface and consists of an outer multinucleated syncytiotrophoblast layer, which arises from fusion of subjacent mononuclear cytotrophoblasts^3,4^. Trophoblast fusion is hence critical for a successful pregnancy as the syncytiotrophoblast regulates transport between mother and fetus, and dysregulated fusion is associated with abnormal placentation that can lead to significant obstetric complications such as pre-eclampsia and intrauterine growth restrictions^5^. While several models exist with which to study placental transport ^6,7^ including recent advanced ‘on-a-chip’ devices^8–13^, creating perfectly fused syncytial sheets remains challenging in culture. While novel pluripotent stem cell models do provide one avenue to enhance fusion efficiencies^14–16^, developing a better understanding of the fundamental factors influencing syncytialization could further improve upon these advanced models, and enhance our understanding of the factors driving disease progression during pregnancy.

The chorionic villus begins developing as a small bud and progresses towards a branched terminal villi encapsulated within a syncytialized trophoblast sheath, through a process of repeated budding morphogenesis^17,18^. Syncytin, cytokines, proteases, growth factors and oxygen levels play a well-established role in successful fusion^19–21^. Conversely, the biochemical factors encountered during diseases including intrauterine growth restriction, gestational diabetes, and pre-eclampsia are known to disrupt fusion and impair villous formation^22^. While the effects of these cues on fusion processes have been well-studied, the impact of biomechanical factors on fusion, particularly those that arise during villous development, remains less clear.

Mechanical forces arising from budding morphogenesis have been established to play an important role in directing cell function in the gut^23^ and lungs^24,25^ for example; and are known to affect fusion in systems such as muscle^26^ and bone^27^. Such stress patterns can spontaneously arise during 3D tissue morphogenesis^28–30^ to directly impact cell function^31^. In placental models specifically, endogenous mechanical stress patterns have been computationally predicted to arise from trophoblast proliferation and fusion^32^, and such endogenous stresses have been experimentally shown to affect trophoblast fusion via models that tune substrate stiffness and colony sizes^33–35^. Hence, it seems likely that the spatially dynamic biomechanical stress profiles that arise during villous development could play an important role in the fusion process.

In this work, we ask whether fusion is associated with altered biomechanical stresses and show that sites of fusion in a trophoblast model of monolayer fusion are accompanied by spatial patterns of mechanical stress consistent with those expected from the development of bud-like projections. We then recreate these patterns of dynamic stress and demonstrate that the application of externally generated, spatially directed compressive bud-like stress enhances fusion efficiency. We extend these findings to a 3D spheroid culture model, and demonstrate that low levels of compressive stress can enhance fusion even without a biochemical induction method of forskolin^36^.

## Results

### Increased compressive radial stresses arise at sites of induced fusion in culture

While the mechanical impact of processes such as proliferation and apoptosis have previously been understood in biological systems, the mechanical features that accompany a trophoblast fusion event have not been previously studied. Here, using traction force microscopy (TFM), we aimed to quantitatively measure the stresses associated with stochastic fusion events observed after forskolin-induction in a classical choriocarcinoma cell model system of trophoblast fusion (Figs. 1A and 2A). A live monolayer of BeWo cells was imaged under forskolin stimulation, and assessed for fusion by identifying those regions positive for Syndecan-1, a marker for successful fusion^37^. In this way, regions exhibiting fusion can be distinguished from those that remain mononucleated, and the history of mechanical stresses in those regions can be quantified.

**Figure 1.**
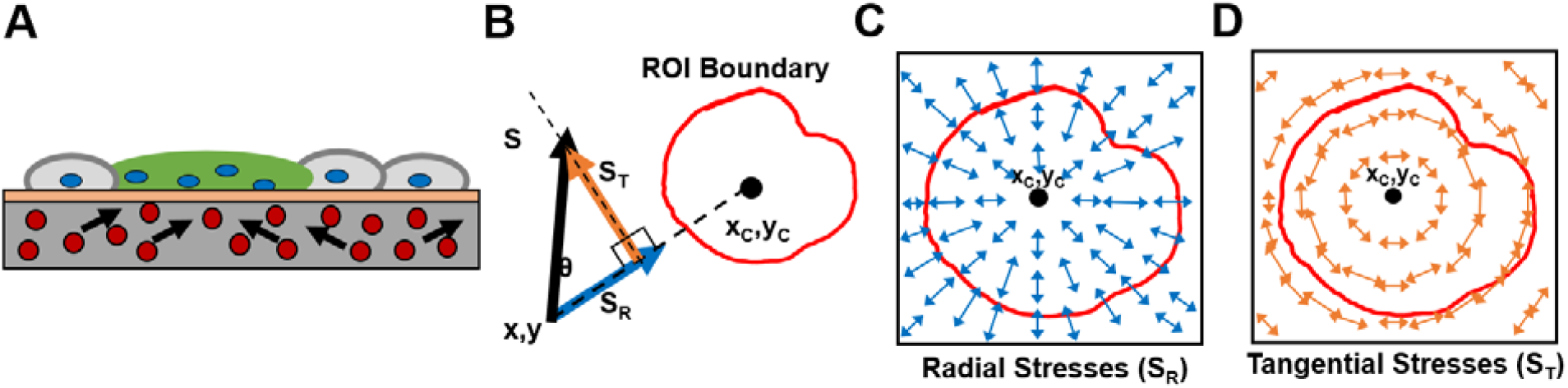
Techniques used for Traction Force Microscopy (TFM) analysis. (A) Schematic illustrating the TFM experiment performed with sheets of fused and non-fused BeWo cells on a polyacrylamide gel. Arrows indicate the change in position of fluorescent beads that is used to calculate stresses. (B) Stress vectors *S* were decomposed into radial *S*_*R*_ (blue) and tangential *S*_*T*_ (orange) components based on the areal centroid (x_C_,y_C_) of a fused region. (C,D) Schematic illustrating an idealized stress field of all the correspondingly resolved radial and tangential stress vectors, respectively within and around a region of interest (ROI).

**Figure 2.**
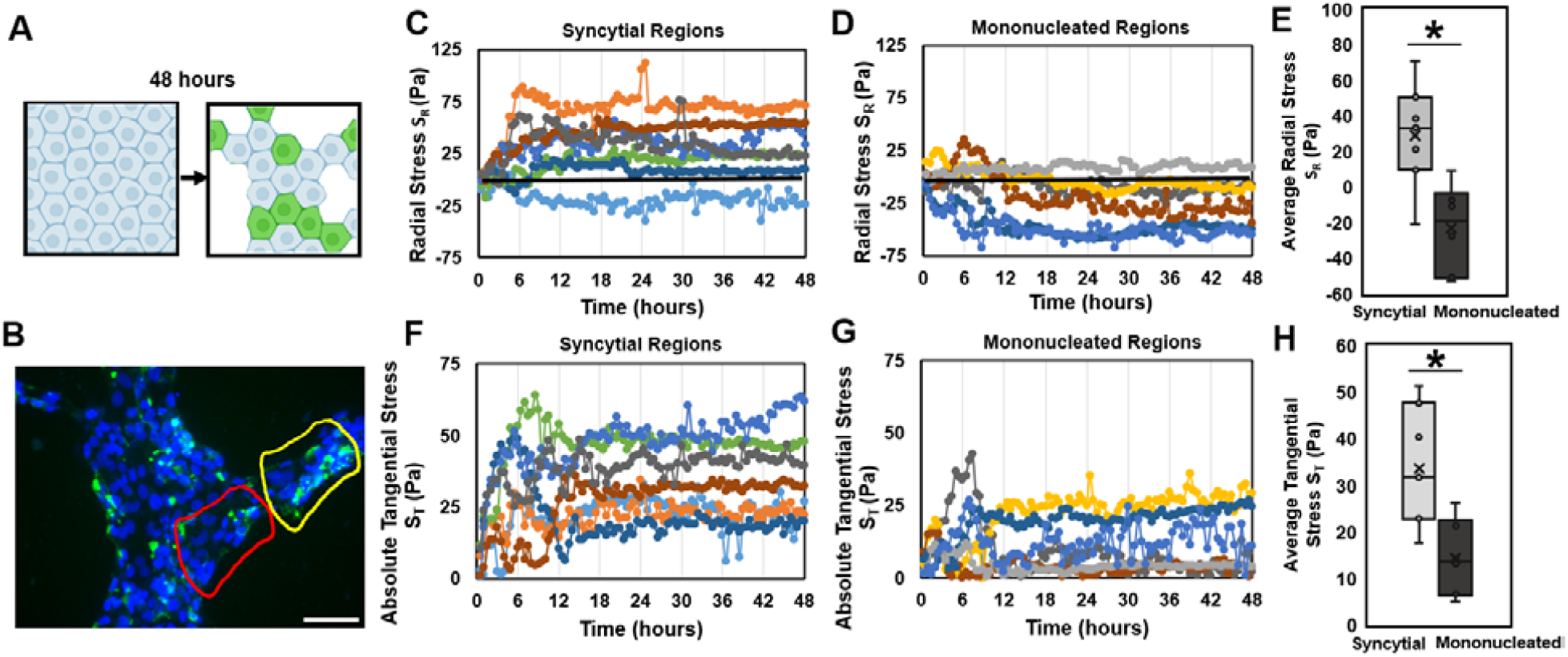
Traction Force Microscopy (TFM) performed on fusion induced BeWo monolayers. (A) Schematic depicting a sheet of cells undergoing fusion over a period of 48 hours; fused regions represented in green. (B) Representative fluorescent image of a sheet of BeWo cells, showing a fused syncytial region (yellow outline) and a non-fused mononucleated region (red outline). Scale bar is 100 μm; green: Syndecan-1, blue: nuclei. (C,D) Radial traction stresses and (F,G) tangential traction stresses over 48 hours, for both syncytial and mononucleated regions, respectively. Each line represents one tracked region. (E) Radial traction stresses and (H) tangential stresses, averaged from hours 12 to 48 for both syncytial and mononucleated regions. (Data presented as box plot distributions; n = 6-7; * *p* < 0.05 by independent Student’s t-test).

As expected, we observed stochastic appearance of small fused (syncytial) patches across the fenestrated monolayer, amidst non-fused (mononucleated) regions (Fig. 2B; overall fusion efficiency of 38 ± 3% across 3 independent samples). We then considered stresses encountered during the prior 48 hours around syncytial and mononucleated areas, and quantified the radial and tangential components of these stresses in relation to the centroid of the syncytial or mononucleated patch (Figs. 1B-D). This approach allowed us to quantify compression and tension in the radial (inwards/outwards) direction (radial stresses or S_R_) as well as circumferential stresses around the edges (tangential stresses or S_T_).

Changes in both radial and tangential traction stresses were observed within ∼6 to 12 hours after induction of fusion and then stayed relatively constant for the duration of the experiment (Figs. 2C, D; F, G). To quantify the mechanical stress levels without capturing this transition period, stresses from 12-48 hours were averaged for comparison purposes (Fig. 2E, H). A statistically significant increase in compressive radial stresses (directed into the region of interest) was observed around syncytialized regions, but not in mononucleated sites of equivalent area (Fig. 2E). Similarly, an increase in tangential stresses (directed around the region of interest) was observed (Fig. 2F-H) for syncytialized regions. This pattern of biaxial compression towards a radial region matches the stresses predicted to arise during villous bud formation due to local proliferation of cells^17,34^, suggesting that the process of fusion itself may also contribute to sculpting the villous bud *in situ*. However, whether these forces are merely correlated with, or a necessary component of the fusion process remains unclear and can only be addressed by recreating these dynamic mechanical stresses in culture.

### Externally applied uniaxial compression does not affect fusion efficiency

To externally apply dynamic mechanical stresses to cells in monolayer culture, we utilized a commercially available elastomeric platform (CellScale MechanoCulture FX) to apply an externally defined strain-based deformation over time. The substrate strains are then expected to apply stresses to the adherent cells, which can then respond and remodel in response to the applied load. We first tested external uniaxial compressive strains, which should apply an appropriate load without matching the spatial patterns of stress observed in the stochastic fusion stress measurement experiments. A loading profile of 10% compression was selected based on the displacements observed in the traction force microscopy experiments; and applied in steps over two days (Fig. 3A, B). As negligible fusion was observed for BeWo cells in standard media, we added forskolin to all cultures for these experiments to compare the relative change in fusion efficiency due to compression. Interestingly, we found that uniaxial compression did not have any effect of fusion efficiency when compared against a static control (Fig. 3C-3E). This would suggest that compression along a single axis is not sufficient to induce fusion, perhaps due to cell reorganization along other axes to relieve unidirectional stresses. If true, this would further suggest that the spatial pattern of force is a key parameter in achieving high levels of fusion in this model.

**Figure 3.**
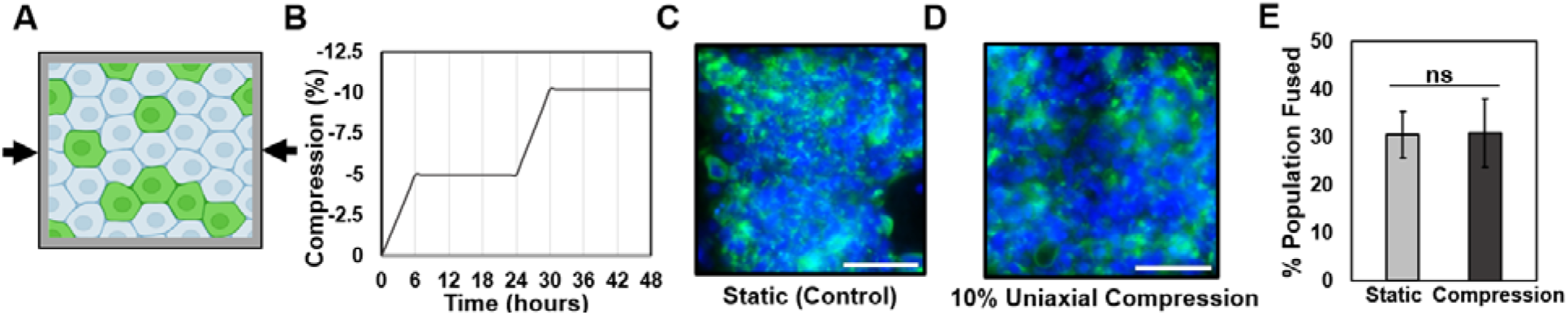
Effect of uniaxial compression on fusion in BeWo cells, induced with forskolin (24 μM) for 2 days. A) Schematic illustrating the 10% uniaxial compression experiment performed along with a graph (B) showing the actual regimen over 48 hours. C) Fluorescent images of fused BeWo cells on a static control and D) subjected to 10% uniaxial compression over 2 days. Scale bar is 100 μm; green: Syndecan-1, blue: nuclei. E) Quantification of fusion efficiency. (Data presented as mean ± standard deviation; n = 5 independent wells, ns-not significant, *p* = 0.916 by independent Student’s t-test).

### Equibiaxial compression increases fusion efficiency whereas equibiaxial tension does not

In order to apply mechanical compression and tension along both the radial and circumferential directions of a circular elastomeric cell culture substrate (Fig. 4A), we used a custom iris-like equibiaxial stretching system^38^. To control for possible confounding effects of substrate stiffness in these experiments, we fabricated a stiffness-matched custom silicone mixture in a petri dish to serve as a static control. No statistically significant differences in mechanical stiffness were observed between the commercially prefabricated stretch dishes and cast silicone elastomer static petri dish samples (Supplementary Fig. S1). We have also previously shown that substrate stiffnesses greater than 50 kPa induce similar morphological phenotypes in BeWo cells as hard tissue culture plastic^33^. Together, this data demonstrates that the control conditions are suitably stiffness-matched so as to isolate the effects of externally applied mechanical stress on the cell cultures.

**Figure 4.**
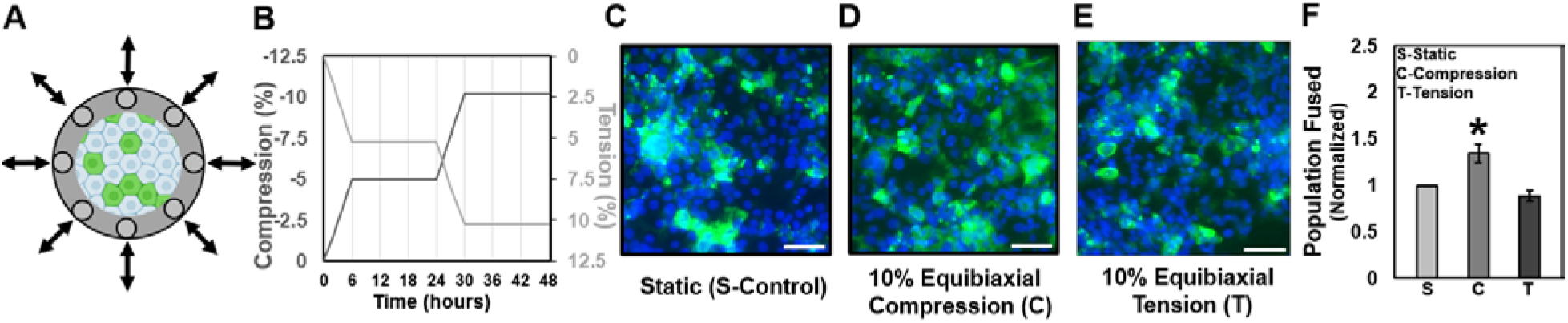
Effect of equibiaxial compression and tension on fusion in BeWo cells, induced with forskolin (24 μM) for 2 days. (A) Schematic illustrating the 10% equibiaxial compression/tension experiment performed along with a graph (B) showing the actual regimen over 48 hours. (C) Fluorescent images of fused BeWo cells on a static control, (D) subjected to 10% equibiaxial compression, and (E) subjected to 10% equibiaxial tension over 2 days. Scale bar is 100 μm; green: Syndecan-1, blue: nuclei. (F) Quantification of the averaged normalized population fusion. (Data presented as mean ± standard deviation; n = 3 independent experiments, * *p* < 0.05 by one-way ANOVA).

Similar to the uniaxial compression experiment, BeWo cells were subjected to 10% equibiaxial strains in either compression or tension over 2 days (Fig. 4B). A statistically significant increase in fusion over static conditions was observed only when cultured under compression (Fig. 4C-4E, F; Supplementary Table S1). These results confirm that for mechanical stimulation to enhance fusion, the stress fields must recreate both the directional (compressive) and spatial (biaxial) stresses as those predicted computationally^34^ and experimentally observed to arise spontaneously during stochastic fusion in monolayer culture (Fig. 2C, F).

### 3D compression confirms mechanical forces from compression positively influence fusion

To determine whether our findings extend to 3D culture models, we investigated the effect of compression on syncytiotrophoblast formation in a spheroidal BeWo model. Trophoblast organoids have recently emerged as important tools to study various aspects of placental biology including self-assembly, expansion and differentiation^15,16,35,39^. However in these models, fusion takes place at the core of the spheroids which is opposite to the *in vivo* situation^15,16^. We have previously shown that the spheroid core is subjected to compressive stresses generated by a ‘skin’ of tension formed on the spheroid surface^28^. Hence, we wondered whether the state of internal compression present in trophoblast organoids might drive internal syncytialization and sought to manipulate the internal compression in such systems.

To experimentally manipulate compressive stresses in 3D culture, we subjected BeWo spheroids to osmotically induced compression (Fig. 5A). Briefly, long-chain molecules in the cell culture media such as dextran are able to alter the osmotic pressure of a solution. Biological objects such as spheroids permit water transport but prevent dextran movement. This establishes an osmotic pressure gradient across the tissue boundary, through which the tissue undergoes compression in an isotropic stress field. This has been previously shown to significantly affect biological systems through processes such as reduced proliferation^40^ or Wnt-mediated stem cell renewal^41^. This technique has the advantage of generating mechanical stresses without complex equipment or structures that is often required for physical manipulation of 3D cultures.

**Figure 5.**
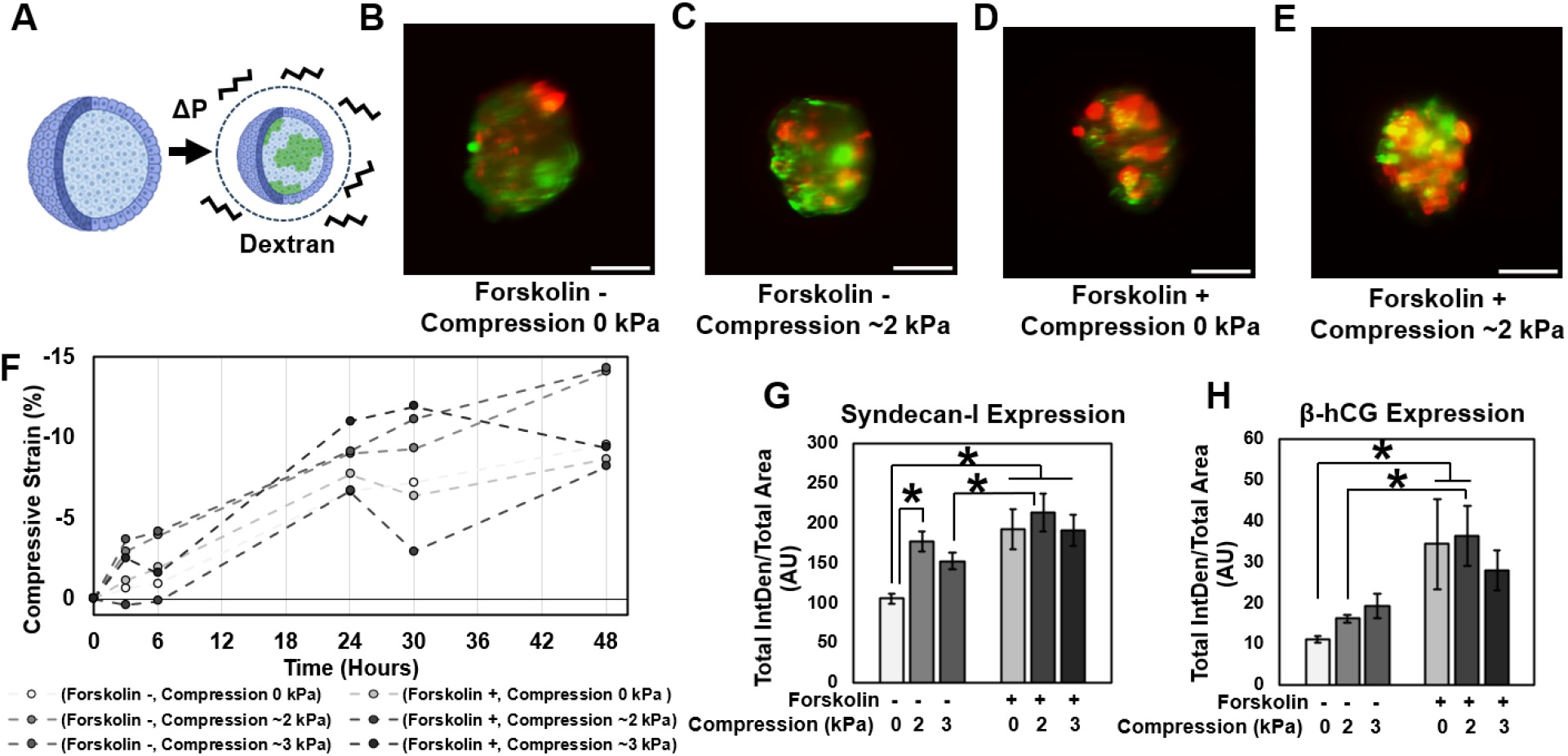
Effect of 3D compression on fusion in BeWo spheroids, induced with (+)/without (-) forskolin (24 μM) and/or dextran for 2 days. (A) Schematic illustrating the 3D compression experiment performed using dextran molecules. (B-E) Representative fluorescent images of a slice of BeWo cells from spheroids subjected to various conditions. Scale bar is 100 μm; green: syndecan-1, red: β-hCG. (F) Trend of quantified compressive strain for spheroids subjected to various conditions at several time points over 48 hours. (G,H) Quantification of fusion using Syndecan-1 and β-hCG markers. (Data presented as mean ± standard deviation; n = 3 spheroids per condition, ^*^ p < 0.05 by two-way ANOVA with post-hoc Tukey’s test)

To quantify the degree of compression generated by a known osmolar imbalance, we first determined the compressive strain exhibited by a polyacrylamide hydrogel of known stiffness (G = 5.15 ± 0.34 kPa, as assessed by shear rheometry) in response to 25 and 50 mg/mL of dextran. Finite element modelling was then used to determine the stresses required to achieve these deformations (1.85 ± 0.67 kPa or ∼2 kPa, and 2.86 ± 2.50 kPa or ∼3 kPa for 25 and 50 mg/mL, respectively). These values are consistent with other studies^42^.

BeWo spheroids were formed in polyacrylamide micropockets^43^ and maintained in culture for two days with and without both forskolin and externally applied compression (Fig. 5B-5E). In all cases, spheroidal tissues exhibited some degree of additional compaction over two days, likely due to continued internal remodelling^28^. However, this compaction was markedly increased in spheroids subjected to dextran compression without forskolin treatment (Fig. 5F). Analysis of nuclear shape in immunostained sections of these tissues indicated that all nuclei were intact and normal (Supplementary Fig. S2), confirming that compression was achieved without inducing apoptosis or necrosis within the spheroidal tissue.

We evaluated fusion efficiency via whole-mount light sheet fluorescent imaging (Fig. 5G-5H) and confirmed our findings with immunostained tissue sections (Supplemental Fig. S2). While the least fusion was observed in the condition without compression or forskolin, we noted that BeWo cells did exhibit some degree of fusion even without forskolin induction, in contrast with our studies of fusion in monolayers which required forskolin. We also noted that 2 kPa of compression was sufficient to significantly increase fusion in standard cultures. Furthermore, at these levels of compression, we achieved fusion efficiencies statistically similar to those achieved via forskolin induction. Surprisingly, while intracellular levels of β-hCG did increase significantly with both compression and forskolin-induction, they did not match the levels of fusion inferred by Syndecan-I expression, consistent with other reports suggesting that intracellular β-hCG levels do not scale with fusion efficiency ^33,44^. This data does however confirm that compression affects fusion in 3D, even in the absence of forskolin induction.

## Discussion

While mechanical forces are now well-established to play an important role in directing cell function^45^, the specific relationships between forces arising during development and the corresponding specification of cell behaviour are highly diverse across organ systems. Placental villous tree formation is a multifaceted and dynamic process, and understanding the precise mechanical contributions and impact of the various processes occurring during villous morphogenesis remains challenging. Here, we demonstrate that the process of fusion itself is associated with specific patterns of local stress at the site of fusion. These findings are consistent with other studies of cell fusion in muscles^46^ in which the highest contractile stresses were observed at the tips of cells undergoing differentiation. Interestingly, the patterns of stress observed in trophoblast fusion are similar to those that would drive morphogenetic budding in the placental bed. We then demonstrate that applying qualitatively similar patterns of external compressive stress can enhance cell fusion efficiency, in both 2D and 3D culture models. These results together suggest that a positive feedback loop exists between trophoblast fusion and mechanical remodelling that act together to sculpt a villous tree structure with a continuous syncytialized monolayer.

We also demonstrate that in 3D cultures, externally applied low levels of compressive stresses can potentially achieve similar levels of fusion as via chemical induction with forskolin. Although BeWo choriocarcinoma cells are a frequently used model for studies of trophoblast fusion, non-physiological chemical induction is often considered essential for fusion in this model. Speculatively, our findings therefore could suggest that downstream effects of chemical stimulation may be physically responsible for fusion. For example, forskolin induction increases production of cyclic adenosine monophosphate (cAMP) which is also known to regulate mechanical contractility in the heart^36,47^, and might establish pro-fusion mechanical conditions. This suggests the intriguing possibility that recently-developed trophoblast stem cell models ^14,48^ that exhibit higher frequencies of spontaneous fusion may present distinct mechanical behaviours and architectures that make fusion more efficient.

Our study does have some significant technical considerations that limit interpretation of results. Specifically, quantitatively recreating the stress patterns measured via TFM and predicted via existing computational models cannot be achieved with current commercially available systems. 2D monolayer mechanical stimulation systems are strain-controlled, rather than stress-controlled, and rely on transfer of deformation patterns from the elastomeric culture substrate to the adherent cells. Conversely, osmotic compression of 3D systems is stress-controlled, but cannot create specific spatially defined deformation patterns. Hence, although our findings correlate well with each other and are conceptually unified, establishing quantitative stress responses could be an important next step for this work. This would require novel technological developments in cell mechanobiology stimulation platforms, perhaps using 3D engineered hydrogel model systems to apply local stresses to living tissue^49^.

More broadly, this specific study does have conceptual limitations that should be considered carefully. First, all experiments were performed with the BeWo choriocarcinoma cell line, which although a well-established model to study fusion processes specifically, may not accurately capture other aspects of placental biology. This is appropriate for a first study because we focus explicitly on establishing external mechanical stresses as fundamental drivers of fusion, but extending this work towards other trophoblast stem cell types would be an important next step. Second, the 2D experiments performed here examine fusion between adjacent cells in culture. Typically, maintenance of the syncytiotrophoblast requires fusion through the basal surface of an established syncytium, which is experimentally quite challenging to recreate. While we can conclude that *in vitro* fusion is impacted by mechanics, whether this translates directly to *in vivo* contexts such as syncytial maintenance or to primary syncytialization remains unclear. Finally, establishing causative mechanistic relationships between external mechanical stresses and fusion is particularly challenging. Unlike molecular systems, “external mechanics” cannot be specifically and precisely inhibited or targeted without affecting a wide variety of cellular processes. Nevertheless, establishing the mechanosensitive pathways that affect fusion would be an important next step in identifying actionable mechanisms to manipulate fusion as needed.

In summary, our results demonstrate that directed external mechanical stresses in the form of compression, such as those that would arise when a small patch of cells fuse, might be an important factor in efficient fusion *in vitro*. We also explicitly show this in spheroids where greater fusion was observed under low compressive stresses even in the absence of forskolin, demonstrating that mechanical stimulation is likely a potent stimulator of fusion and can be utilized as a tool to impact *in vitro* levels of fusion. Broadly, this work provides a better mechanobiological understanding of trophoblast fusion as a fundamental biological process, which can ultimately be leveraged to improve fusion efficiency in *in vitro* models and potentially be used as a novel therapeutic target for placental dysfunction *in vivo*.

## Methods

Unless otherwise stated, all cell culture materials and supplies were purchased from Fisher Scientific (Ottawa, ON) and chemicals from Sigma Aldrich (Oakville, ON).

### Cell Culture - Monolayers

BeWo human placental choriocarcinoma cells (ATCC; CCL-98) between passages 20-25 (for monolayers) or passages 4-6 (for spheroids) were cultured in 10% foetal bovine serum (FBS) and 1% antibiotic-antimycotic in Dulbecco’s Modified Eagle Media (DMEM). BeWo cells were seeded at ∼120,000 cells/cm^2^ to form a continuous monolayer sheet of cells. Media was supplemented with 24 μM forskolin (CAS: 66575-29-9) to induce fusion, with media replaced every 24 hours. For traction force microscopy (TFM) experiments, an excess volume of media was used to avoid disrupting the automated imaging for a media exchange.

### Cell Culture - Spheroids

A custom stamp block with a 1 mm cone at the surface was adapted from a previous technology^43^, and designed to fit a 48 well plate using Fusion 360 (Autodesk) and printed in PR57-K black prototyping resin (Colorado Photopolymer Solutions) using the Autodesk Ember DLP 3D printer. To create micropockets, Loctite AA3525 was used as an adhesive to attach polyacrylamide gels to the bottom of the well plate^50^. The adhesive was incubated for 1.5 hours in RO water to limit toxicity, after which a polyacrylamide mixture corresponding to 25.6 kPa shear stiffness^51^ was added to the bottom of the well and covered immediately with the stamp block. After curing for 8 minutes, the block was removed and left in PBS solution containing 1% antibiotic-antimycotic for at least 3 days and with the solution changed every 24 hours to remove any residual chemicals that might be harmful to cells. On the day of experimentation, microwells were exposed to 365 nm UV light for 1-2 hours and the incubated in complete culture media. 50,000 cells were pipetted into the well containing the micropocket and the well plate was centrifuged at 400 RCF for 10 minutes, after which the well plate was incubated at 37 °C for 1-2 days to enable spheroids to form.

### Traction force microscopy

We have previously determined polyacrylamide hydrogel formulations that result in gels with stiffnesses that mimic native healthy placental tissue mechanical properties^33,52^. For TFM experiments, prepolymer solutions of the 3900 kPa native placental stiffness containing 0.5% volume of 0.5 μm diameter carboxylate-modified fluorescent beads in PBS (FluoSpheres, Invitrogen, Catalog: F8812) were prepared as previously described^34^. To functionalize the gel surface, the gels were activated twice with 0.1 mg/ml of the photoactivable bifunctional crosslinker N-sulfosuccinimidyl-6-[4’-azido-2’-nitrophenylamino] hexanoate (sulfo-SANPAH, ProteoChem); and incubated with an excess amount of 80-100 μl of bovine Collagen ı (0.1 mg/ml in phosphate buffered saline or PBS, Life Technologies) on a parafilm sheet at 4 °C overnight. Such TFM substrates prepared on 12 mm glass coverslips were attached to the bottom of a 24 well plate using a drop of cured polydimethylsiloxane (PDMS, 10:1 pre-polymer: curing agent, Dow Sylgard 184) and exposed to 365 nm UV light for at least 45 minutes to facilitate quick attachment and to prevent detachment of gels, after which gels were washed in sterile PBS and immediately used for experiments.

TFM substrates were incubated in complete DMEM for 2 hours at 37 °C and then with BeWo cells, which were allowed to attach and form a uniform monolayer over 24 hours. To capture the stressed conditions required for the traction force analysis, red fluorescent images of the uppermost layer of embedded beads were captured along with phase-contrast images of the selected regions every 30 minutes using an automated fluorescent microscope (20× objective, EVOS m7000, ThermoFisher Scientific) for a total time of 48 hours. The first time point at t=0 hours was used as the reference to obtain the relative traction forces over the imaging period. After 48 hours, cells were fixed and immunostained, then the same regions were re-imaged to identify fused syncytial regions and non-fused mononucleated regions.

### TFM image analysis

To analyse the TFM image sets, template alignment, particle image velocimetry (PIV), and Fourier transform traction cytometry (FTTC) ImageJ plugins were used for the TFM analysis, following previous protocols^53^. Briefly, both stressed and relaxed fluorescent bead images were combined into a stack and aligned to account for experimental drift. The bead displacements were estimated by PIV (advanced) following an iterative procedure where the interrogation window was made progressively smaller (128×128 pixels, 64×64 pixels, 48×48 pixels; correlation threshold: 0.6) to produce an ultimate displacement field grid of ∼14.8 μm x 14.8 μm. Traction force fields could then be reconstructed with the FTTC plugin using values: pixel=0.3086 μm, Poisson’s ratio = 0.457, and Young’s Modulus = 3900 Pa, along with default values for remaining inputs. This was repeated for each time point over the entire 48 hours using a custom ImageJ macro.

Identified fused syncytial regions as evidenced by presence of Syndecan-1^37^ on stained images at t=48 hours were manually identified in ImageJ and enlarged by 80 pixels to include 1-2 cells outside the region of fusion to include their influence on the stress patterns. A corresponding region of same area but for a non-fused mononucleated region was also selected and all regions of interest (ROIs) were utilized by a custom MATLAB code to project the traction stresses of those ROIs towards the centre as well as tangentially (perpendicular, anti-clockwise), and to calculate the average radial and average absolute tangential stresses within the ROIs at each 30 minutes time point over the entire 48 hours. For quantification purposes, radial stress values from t=12 to 48 hours were further averaged to be used for comparison between syncytial vs. mononucleated regions.

### Uniaxial compression stimulation experiments

Uniaxial compression experiments were performed with a commercial mechanical stimulation platform (CellScale MechanoCulture FX) using a 24 well silicone membrane platform. The silicone platform was pre-stretched to 10% uniaxial strain, coated with 0.1 mg/ml bovine Collagen I by incubating at 37 °C for 2 hours, seeded with cells in the stretched condition and left undisturbed for 24 hours to form a sheet. Similar steps were performed on a separate static 24 well silicone membrane platform to be used as a control. A mechanical stimulation regimen consisting of 5% uniaxial compression over 6 hours and no movement for 18 hours was repeated twice for a total of 2 days, so that an overall 10% uniaxial compressive strain could be achieved. Media containing forskolin was changed daily. After 2 days of stimulation, cells were fixed, immunostained, and imaged.

### Equibiaxial compression and tension stimulation experiments

Both equibiaxial compression and tension experiments were performed using a custom iris-like stretchable device system that has been previously described in multiple stretching applications^38,54^. In order to establish a static control to compare against the commercial pre-fabricated stretchable dishes used for the stimulation experiments, a silicone membrane with similar properties to that of the stretchable dishes was replicated in a 35 mm petri dish by mixing equal parts of Silicone Elastomer Parts A and B (Elkem LSR-4305 Elastomer, Product Code: A-221-05; Factor II, Inc.) and cured in an oven at 70 °C for 2 hours after degassing. The Young’s modulus of both the static control petri dishes and stretchable dishes were verified using rheometry (see Supplementary Methods S1). Both the stretchable dishes and the static control petri dishes containing silicone were washed with double distilled water (ddH_2_O) and treated with 30% sulfuric acid for 15 minutes at room temperature. Both platforms were again rinsed with ddH_2_O and then treated with 1% (3-aminopropyl) triethoxysilane (APTES) for 2 hours in a 70 °C oven. Both platforms were again rinsed with ddH_2_O and stored in a 4 °C refrigerator for up to a month prior to use in experiments.

On the day of experimentation, both platforms were treated with 1% glutaraldehyde solution for 15 minutes at room temperature after which both platforms were rinsed with ddH_2_O, sprayed with 70% ethanol, and taken into a biological safety cabinet for sterility with subsequent steps. The stretchable dish was connected to a computer to be pre-stretched using a custom LabVIEW based software to the required base minimum area (for compression experiments) or pre-stretched to 10% strain from the base minimum area (for tension experiments). Both platforms were coated with 0.05 mg/ml Collagen I by incubating at 37 °C for 2 hours. After rinsing with sterile PBS, cells were seeded in both platforms and left undisturbed up to 24 hours to form a monolayer. A mechanical stimulation regimen consisting of 5% equibiaxial compression (or tension) over 6 hours and no movement for 18 hours per day was repeated twice for a total of 2 days, so that an overall 10% equibiaxial compressive (or tensile) strain could be achieved. Media containing forskolin was changed daily. At the end of 2 days, cells were fixed, the bottom of stretchable dishes were cut out with a scalpel, and then immunostained and imaged. Experiments were repeated for 3 independent stretchable dishes and static petri dishes with results normalized to the corresponding control for comparison.

### Immunostaining

At the end of uniaxial compression, equibiaxial compression/tension and TFM experiments, 8% (w/v) formalin in PBS was added to samples containing 50% remaining media to fix for 12-15 minutes at room temperature. Samples were then rinsed in PBS, permeabilized with 0.1% (v/v) Triton X-100 in PBS for 12-15 minutes, washed in PBS, blocked with 2.6% (v/v) goat serum, and incubated overnight at 4 °C with primary rabbit anti-Syndecan-1 antibody (HPA006185, 1:200 dilution) for fusion analysis. Secondary staining was performed with goat-anti mouse IgG H&L antibody (Alexa Fluor® 594 red, 1:1000 dilution) and goat anti-rabbit IgG H&L antibody (Alexa Fluor® 488 green, 1:1000 dilution, Invitrogen) for 3 hours, and counterstained with Hoechst 33258 (5 μg/ml) for 2 hours. For whole spheroids with all steps performed overnight at 4 °C, samples were fixed with 4% (w/v) formalin in PBS, permeabilized and blocked together in a buffer solution containing 0.5% Triton X-100 and 5% goat serum in PBS (v/v), primary stained with goat anti-Syndecan-1 antibody and mouse anti-human choriogonadotropin (β-hCG) antibody (#14-6508-82, 1:500 dilution in the buffer solution, ThermoFisher), and secondary stained as previously described. All samples were kept at 4 °C until image acquisition.

### Osmotic compression of 3D cultures

Spheroids were used for fusion-compression experiments under the combinatorial conditions of with/without forskolin and with/without dextran, where dextran (Mw = 500 kDa, Dextran Products; Polydex Pharmaceutical) was added separately as 25 mg or 50 mg of dextran per ml of complete DMEM media. Media was changed every 24 hours and timelapse images at time points t = 0, 3, 6, 24, 30 and 48 hours were acquired to assess compaction though compressive strains. After 48 hours, samples were washed with PBS and then fixed overnight for immunostaining or histology to confirm findings (see Supplementary Methods S2).

### COMSOL Modelling: Determination of spheroid compression due to dextran

The isotropic compression applied on the spheroids due to the concentration of dextran used was calculated using a model similar to previous work^28^. In brief, disk-shaped polyacrylamide gels were fabricated by casting polyacrylamide prepolymer gel solution (18.75% v/v 40% Acrylamide, 2.70% v/v 2% Bisacrylamide, 68.40% v/v PBS, 0.15% v/v TEMED and 10.0% v/v APS) into 12 mm molds under a constant nitrogen gas flow for 15 minutes. After gelation, disc gels were washed with PBS 3 times and stored at 4 °C to allow the gels to swell. After 7 days, the rheological properties of disc gels were measured through rheometry (see Supplementary Methods S1). We have previously demonstrated that this dextran chain length formulation does not penetrate polyacrylamide gels ^28^. The change in size of the gels due to dextran induced osmotic compression was measured using dextran solutions with concentrations of 25 mg/ml and 50 mg/ml in PBS. To do so, images of the disc bulk gel were acquired using the EVOS m7000 microscope at 4X magnification before and 30 minutes after addition of Dextran solution at 37 °C, and the corresponding change in diameter was calculated in ImageJ (NIH).

To quantify the compressive forces generated by the dextran, an inverse finite element simulation was made in COMSOL Multiphysics 6.0 (Comsol Inc., Burlington, MA, USA). Disk shaped geometries were generated in a 2D axisymmetric model, and a pressure boundary condition was set on the top and side while a roller condition was placed on the bottom surface. A free triangular mesh was generated with predefined extremely fine element size giving an average element quality above 0.9. Disks were prescribed linearly elastic properties with a Young’s modulus measured by rheometry (see Supplementary Methods S1) and a Poisson ratio estimated at 0.457 from literature^55^. Using the experimentally measured pre- and post-compression gel sizes, the applied pressure was determined.

### Microscopy

All samples of equibiaxial compression or tension experiments were imaged on an inverted fluorescent Olympus microscope (Olympus, IX73) outfitted with an sCMOS Flash 4.0 Camera and Metamorph software (version 7.8.13.0) for brightfield and corresponding fluorescence filters (blue for Hoechst 33258, red for E-Cadherin and green for Syndecan-1). For TFM experiments, the uniaxial compression experiment, spheroid compression time lapse experiment and spheroid histology sections, samples were images on an inverted EVOS m7000 microscope for brightfield and corresponding fluorescence filters (blue for Hoechst 33258, green for Syndecan-1 and red for fluorescent beads or E-cadherin). Images were acquired at random locations at 20X magnification for stimulation experiments and spheroid sections, at fixed positions at 20X magnification stored in memory over 48 hours at 30-minute intervals for TFM experiments, and per well at 4X magnification over 48 hours at time points t = 0, 3, 6, 24, 30 and 48 hours for the spheroid compression time lapse experiment.

Whole spheroid samples were imaged on a customized SCAPE 2.0 light sheet microscopy system^56^ in a top-down format, outfitted with a water dipping objective lens (20X 1.0 NA) and an Andor Zyla 4.2+ CMOS camera. Samples were individually encased in 1% agarose (UltraPure™ Agarose, Invitrogen) in a 35 mm petri dish prior to imaging. Slices of the entire sample were obtained at 10 μm spacing under 9.3X magnification, using a 488 nm excitation laser with 100 ms exposure and 10 mW power at source settings (green for Syndecan-1) and a 561 nm excitation laser with 100 ms exposure and 8 mW power at source settings (red for β-hCG).

### Image Analysis

All images were analysed in FIJI ImageJ (NIH). For fusion analysis, a colony containing 2 or more cells sharing the same membrane boundary and/or strongly expressing Syndecan-1 was considered to be fused. The total number of nuclei was counted either manually or using the ParticleSizer plugin in ImageJ^57^ to calculate fusion efficiency as described below, based on similar techniques previously reported^34,37,58^:

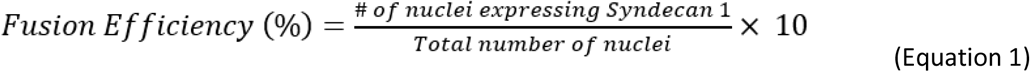

For fusion analysis in spheroids, the brightness and contrast of all slices for all conditions for both Syndecan-1 and β-hCG were independently fixed to a specific value and the IntDens values of the spheroid in all slices of the stack were obtained. The area of the spheroid in all slices of the stack was obtained through thresholding and using the %Area value for a defined ROI. Fusion levels were calculated as follows:

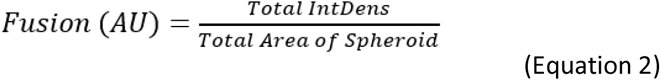

Compressive strains in timelapse images of spheroids subjected to forskolin and/or dextran compression was calculated by finding the effective change in diameter of the spheroid at each time point (assuming a spherical structure) compared to that at t=0 hours. Immunostained sectioned spheroid images were created by overlaying all channels (nuclei, Syndecan-1 and E-cadherin) with brightness and contrast values fixed independently for each channel.

### Statistical Analysis

Statistical analysis was performed using JASP, version 0.16.3^59^. Independent Student’s t-test was used for the uniaxial fusion analysis and comparison of TFM stresses, one-way ANOVA with post-hoc Tukey’s test for pairwise comparisons was performed for equibiaxial compression and tension fusion analyses whereas two-way ANOVA with post-hoc Tukey’s test for pairwise comparisons was performed for spheroid fusion analyses. In all cases, p ≤ 0.05 was considered statistically significant.

### Schematics

Schematics for Figures 2-5 were created with BioRender.com.

### Data availability

The custom codes (ImageJ Macro and MATLAB code) utilized for TFM analysis as well as the raw images obtained for each of the experiment described in this work are accessible via the Open Science Foundation at: https://osf.io/23ypd/?view_only=5575a67edb264367af652025a6a4beba

## Acknowledgements

We thank Nikita Kalashnikov for assistance with the preliminary optimization of TFM experimental and analysis procedures, Raymond Tran for providing his MATLAB code that was modified and utilized for the TFM analysis performed here, Nicholas Wong and James Reeves for assistance with operation of the rheometer, Andrei Bocan for assistance with image analysis involved for COMSOL simulations, and Tarek Klaylat and Mustafah Fakih for assistance in preparation of stretchable devices for equibiaxial experiments.

This work was supported by the Fonds de Recherche du Quebec (FRQ) - Nature et technologies (grant no. 205292, to C.M. and C.V.), Natural Sciences and Engineering Research Council of Canada (RGPIN-2015-05512 to C.M and RGPIN-2019-06778 to C.V), and the Canada Research Chair in Advanced Cellular Microenvironments to C.M. Authors gratefully acknowledge personnel funding from the McGill Engineering Doctoral Award (MEDA) and NSERC– Postgraduate Scholarship (PGS-D) Award to P.K.P.

## Author contributions

Experiment design: P.K.P, C.V., C.M. Experiments: P.K.P., C.L., K.A., J.J., S.R., B.E.C., H.L. Manuscript writing: P.K.P., K.A., S.R., B.E.C., D.H.R., C.V., C.M. The final version has been approved by all the authors.

### Competing interests

The authors declare no competing interests.

## Notes

https://osf.io/23ypd/?view_only=5575a67edb264367af652025a6a4beba

### Competing Interest Statement

The authors have declared no competing interest.

