## Supplementary Information for "Directed biomechanical compressive forces enhance fusion efficiency in model placental trophoblast cultures"

**Supplementary Methods and Data**

***S1. Rheometry***

Shear rheological properties were investigated using a rheometer (MCR 302 Anton Paar Modular Compact Rheometer), equipped with an 8 mm diameter parallel plate measuring system geometry. All samples were tested on a heated stage at 37 °C. A thin piece of silicone layer from the 35 mm petridish static control as well as the prefabricated Cellerator devices was cut and loaded at the center of the testing stage with excess sample trimmed prior to testing. An amplitude sweep of increasing shear strain was completed from 0.01% to 100% strain and the corresponding shear stress, storage modulus, and loss modulus were recorded. 3-4 regions of both the Cellerator device or static control petridish were utilized and the average of the storage modulus under 1% - 10% strain across the pieces was taken to be the shear modulus (S) from which the Young’s modulus (E) was calculated using the formula: E=2S(1+ν), where ν is the Poisson’s ration (0.49 for PDMS). Experiments were repeated for three independent Cellerator devices and petri dishes with results averaged for comparison. Similar experiments were repeated on four independent polyacrylamide gels for measuring the rheological properties of bulk gels to be used for COMSOL modelling,

***S2. Histology***

Histological procedures for BeWo spheroids were adapted from a previous protocol established for placental spheroids made with primary cells^1^. In brief, fixed spheroids were removed from the PAA micropocket and individually isolated into a 48 well plate. Samples were dehydrated for 10 minutes in 70% ethanol and then stained with Eosin (0.1% (w/v) solution) for 30 minutes. Samples were then washed in 70% ethanol and then encased in 2% agarose solution (UltraPure^TM^ Agarose, Invitrogen) for ease of handling during subsequent steps. Samples were then inserted into cassettes and then dehydrated with denatured ethanol in the following sequence: twice at 70% for 30 minutes each, followed by 10 minutes each at 80%, 96%, 96%, 100% and 100%. Samples were then incubated twice with Xylene for 30 minutes each to remove the alcohol and then embedded in Histoplast paraffin wax (FisherScientific) twice at 65 °C for 30 minutes each with new wax for each cycle and then finally stored in a 4 °C refrigerator until further use.

Samples were cut into 5 μm sections using a microtome (Leica RM2235), transferred on to a charged glass slide (VistaVision™ HistoBond® Adhesive Microscope Slides, VWR), dried at 37 °C overnight and then stored at 4 °C until further use. On the day of immunostaining, samples were pre-warmed at 60 °C for 1 hour, deparaffinized in Histo-Clear II (VWR) for 10 minutes and then rehydrated in the following sequence: 3 minutes each at 100%, 90%, 80% and 70% ethanol, followed by water and PBS. Samples were then subjected to head induced epitope retrieval in a 10 mM citric acid buffer solution (Trisodium Citrate Dihydrate (Fisher), pH 6.0) using a pressure cooker (Instapot, Amazon) set to a 20-minute cook time under the regular pressure cook setting. Once complete, samples were allowed to cool and renature for 20 minutes, and then subjected to immunostaining.

For spheroid histological sections, samples were rinsed with PBS, and permeabilized with 0.1% Triton X-100 in PBS (PBS-T) for 5 minutes. Regions containing spheroids were marked using a diamond scribe (McMaster-Carr), blocked in a solution containing 10% goat serum + blocking buffer (0.15% (w/v) glycine powder (CAS: 56-40-6) + 0.2% (w/v) bovine serum albumin (BSA) in PBS-T) and then incubated overnight at 4 °C with primary mouse monoclonal anti-E-Cadherin antibody (1:200 dilution) and primary rabbit anti-Syndecan-1 antibody (1:200 dilution). Samples were rinsed in PBS-T and then incubated for 1 hour at room temperature with goat-anti mouse IgG H&L antibody (Alexa Fluor® 594 red, 1:1000 dilution) and goat anti-rabbit IgG H&L antibody (Alexa Fluor® 488 green, 1:1000 dilution) along with Hoechst 33258 (5 µg/ml) in a 1% goat serum + blocking buffer solution. Finally, samples were rinsed with PBS-T, mounted with Fluoromount under a 22x60 mm glass coverslip (FisherScientific), sealed with nail polish (Sally Hansen - Hard as Nails Xtreme Wear, Invisible) and imaged immediately after drying.

**Supplementary Tables**

**Table S1.** Quantification of fusion in equibiaxial compression and tension experiments. Experiments were independently repeated for n=3 times, and a fold change in each case was calculated to normalize against the corresponding static condition prior to comparison.

| **Equibiaxial Experiments; % Population Fused** | | | | | |
| --- | --- | --- | --- | --- | --- |
| **Condition** | **Expt. 1** | **Expt. 2** | **Expt. 3** | **Average** | **StdDev** |
| **Static** | 20.64 | 23.92 | 14.82 | 19.79 | 3.76 |
| **Compression** | 29.83 | 28.94 | 20.47 | 26.42 | 4.21 |
| **Fold Change** | 1.44 | 1.21 | 1.38 | 1.34 | 0.10 |
| **Static** | 19.74 | 24.39 | 17.29 | 20.47 | 2.95 |
| **Tension** | 18.91 | 20.99 | 14.30 | 18.07 | 2.79 |
| **Fold Change** | 0.96 | 0.86 | 0.83 | 0.88 | 0.06 |

**Supplementary Figures**

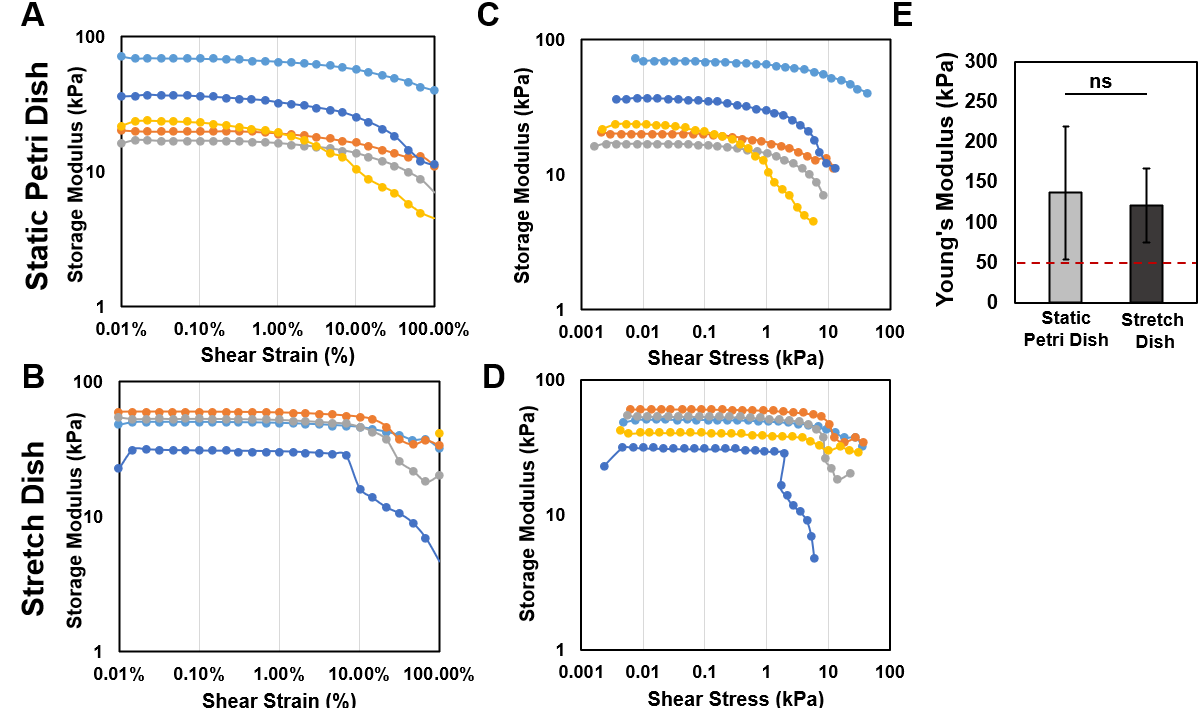

**Figure S1**. Comparison of stiffness between the custom fabricated static control petri dishes and commercially prefabricated stretch devices used for equibiaxial stimulation experiments. A-D) Rheology graphs showing change in storage modulus over increasing shear strains and shear stresses for representative sample sections of both static control petri dishes (A+C) and Cellerator dishes (B+D). E) Quantification of Young’s modulus. RED - Value of Young’s Modulus of previously fabricated polyacrylamide gels at/above which BeWo cells see no differences in stiffness of substrates, taken from Ma *et al.*^2^*.* (Data presented as mean ± standard deviation; n = 3 independent devices each, ns-not significant, *p* = 0.826 by independent Student’s t-test).

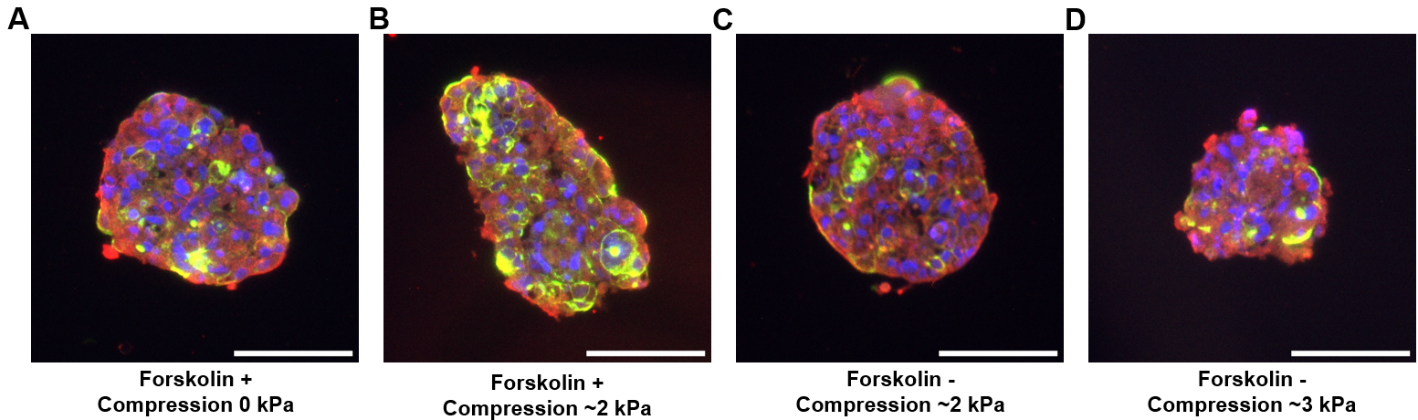

**Figure S2**. Comparison of effect of dextran on nuclei in BeWo spheroids, induced with (+)/without (-) forskolin (24 µM) and/or dextran for 2 days. A-D) Representative fluorescent images of a sectioned slice of BeWo cells from spheroids subjected to various conditions. Scale bar is 100 µm; red: E-cadherin, green: syndecan-1, blue: nuclei (n = 3 spheroids per condition).
